# Range Extension of *Cryptonanus agricolai* (Didelphimorphia, Didelphidae) and First Record in the Atlantic Forest Core

**DOI:** 10.1101/774752

**Authors:** Edú Baptista Guerra, Leonora Pires Costa

## Abstract

According to the Wallacean shortfall, knowledge about the geographic distribution of most species is still incomplete. *Cryptonanus agricolai* (Moojen, 1943) is a didelphid marsupial considered Data Deficient by IUCN, since species records are few and sparse. Although little information is available for the species, it is commonly associated with xeric habitats from Caatinga and open formations of the Cerrado in east-central Brazil. Here we report the first records of *C. agricolai* in the Atlantic Forest core, a new ecoregion of occurrence for the species, based on a recent collected voucher - identified through morphological and molecular analysis - from a Mussununga formation in Reserva Biológica do Córrego do Veado, southeastern Brazil. This record extends the occurrence of the species to more than 1 700 000 km^2^ and lower its altitudinal range limit to 108 m.

**RESUMO:** **Ampliação da distribuição de *Cryptonanus agricolai* (Didelphimorphia, Didelphidae) e primeiro registro no centro da Mata Atlântica.** De acordo com a Lacuna Wallaceana, o conhecimento sobre a distribuição geográfica da maioria das espécies está incompleto. *Cryptonanus agricolai* (Moojen, 1943) é um marsupial didelfídeo classificado pela IUCN na categoria Dados Insuficientes, uma vez que os registros existentes são poucos e esparsos. Embora haja pouca informação disponível para tal espécie, ela é comumente associada a habitats xéricos da Caatinga e formações abertas do Cerrado no centro-leste do Brasil. Aqui relatamos os primeiros registros de *C. agricolai* na Mata Atlântica, notadamente uma nova ecorregião de ocorrência para a espécie, com base em um espécime recentemente coletado – e identificado através de análises morfológicas e moleculares - em formação de Mussununga na Reserva Biológica do Córrego do Veado, sudeste do Brasil. Nossos achados ampliam a ocorrência da espécie para mais de 1 700 000 km^2^ e estabelece novo limite inferior de altitude para 108 m. Palavras-chave. Corredor Central da Mata Atlântica. Lacuna Wallaceana. Marsupial. Mussununga.

## INTRODUCTION

On the way to extend the knowledge of biodiversity, scientists often come across shortfalls on local-scale data. Those shortfalls generate uncertainty in all analyses of biodiversity, compromising the generality and validity of theoretical knowledge and the quality of conservation assessments and actions (Hortal et al. 2015). One of those recognized shortfalls is the Wallacean, which defines that, for the majority of taxa, geographical distributions are still poorly understood and contain many gaps, being frequently inadequate at all scales (Lomolino 2004; Whittaker et al. 2005). Such shortfall is particularly true for tropical species, especially for those hardly trapped and/or not abundant in communities, even though many of them possess high conservation value (Beck et al. 2018).

The Neotropical region is home to one of the richest mammal faunas in the world (Antonelli & Sanmartini 2011; Patterson & Costa 2012). Nevertheless, much of the diversity it houses is still unknown and there are huge sample gaps, even in regions with a long history of occupation and research, as is the case of the Atlantic Forest (Bovendorp et al. 2017). Regarding the knowledge about Neotropical marsupials, within the last years the number of Didelphidae species has increased from 91 species (Gardner 2008) to 103 recognized species (Astúa 2015). However, 15% of these are currently considered as Data Deficient (IUCN 2019). Although research, and consequently, the knowledge on the Neotropical fauna has increased in recent years, gaps in taxonomic and biogeographic knowledge of specious and yet elusive groups, such as the Neotropical small mammals, hinder conservation initiatives. Effective biodiversity conservation requires minimal knowledge about the targets of protection (Brito 2004), making it essential that efforts be devoted to cataloging, quantifying and mapping such biodiversity.

*Cryptonanus* is a didelphid genus that was previously referred as *Gracilinanus*, until be recently validated as a different genus through morphological characters and molecular markers (Voss et al. 2005). Currently, four species of *Cryptonanus* are recognized: *C. agricolai* (Moojen 1943), *C. chacoensis* (Tate 1931), *C. guahybae* (Tate 1931) and *C. unduaviensis* (Tate 1931), which are distributed throughout open habitats in tropical and subtropical ecoregions east of the Andes and south of the Amazon River, including the Caatinga, Chaco, Cerrado and Pampas (Voss et al. 2005; D’Elía & Martínez 2006; Voss & Jansa 2009; Garcia et al. 2010; Quintela et al. 2011).

*Cryptonanus agricolai*, is mostly found in xeric habitats in Caatinga and open formations of the Cerrado biomes in east-central Brazil, from 400 to 760 m (Gardner 2008), but also occurs in contact zones with the Amazon (in northern Mato Grosso state, MN 66318; Bezerra et al. 2009) and the Atlantic Forest limits (seasonal tropical forests of Minas Gerais states; Gardner 2008). Although there is few information available about its locomotion habits, most faunal surveys report captures in pitfall traps, suggesting a ground-dwelling locomotion (Astúa 2015), and probably an insectivore/omnivore diet. The conservation status for such species is assessed as Data Deficient (Carmignotto et al. 2016), since there are few and scattered records over a wide area, and little is known about its habitat requirements, which makes research efforts on its geographical distribution needed.

Here we report the first records of *Cryptonanus agricolai* in the Atlantic Forest core in southeastern Brazil, due to the identification - through morphological and molecular analysis - of a recently collected specimen from state of Espírito Santo, and the recognition of a previously reported *Cryptonanus* sp. (Delciellos et al. 2016) from Rio de Janeiro state as *C. agricolai*. Our findings extend the occurrence of this species to 1 773 398 km^2^ and set its lowest altitudinal range limit to 108 m, in addition to register its presence in an unexpected ecoregion.

## MATERIALS AND METHODS

The Córrego do Veado Biological Reserve comprises a forested area of 2 357 ha in the municipality of Pinheiros, state of Espírito Santo, southeastern Brazil, where secondary vegetation predominantes (Ibama 2000). The surroundings of Córrego do Veado are characterized by anthropic activities, such as cattle ranches and coffee, papaya, eucalyptus and rubber tree plantations (Moscal 2012). On October 17th, 2015, a specimen of didelphid marsupial (LPC1636) was caught in a pitfall trap at Córrego do Veado (18° 19′ S, 40° 07′ W, altitude = 108 m) in an area of Mussununga (i.e. “soft and wet white sand”, in Tupi-Guarani; Meira-Neto et al. 2005), characterized by shrub and bush vegetation that develops in sandy soils (Saporetti-Junior et al. 2012). The individual was collected as voucher and deposited in the Mammal Collection at Universidade Federal do Espírito Santo (UFES-MAM 3075). Liver tissue samples were preserved in 70°GL ethanol and deposited in the Animal Tissue Collection at Universidade Federal do Espírito Santo (UFES-CTA 4116).

Biological information collected for the individual included: reproductive stage, dental age class, body mass (BM, in grams) and external body measurements (in millimeters), such as head-to-body length (HB), tail length (TL), hindfoot length (HL) and ear length (E). Skulls were removed, cleaned, and measured using a digital pachymeter accurate to 0.01 mm. The following cranial and dental measurements were taken from the specimen following Voss et al. (2005): condylobasal length (CBL), nasal breadth (NB), least interorbital constriction (LIB), zygomatic breadth (ZB), palatal length (PL), palatal breadth (PB), maxillary toothrow length (MTR), length of M1 to M4 (LM), and length of M1 to M3 (M1 – M3).

Additionally, 801 bp of the mitochondrial gene cytochrome b (Cyt-b) were used for molecular analyses, due to its potential to distinguish sister species of mammals (e.g. Agrizzi et al. 2012; Bradley & Baker 2001) and also because there are several Cyt-b sequences available for a considerable number of didelphid species in GenBank. Sequence alignment was performed using CLUSTALW algorithm implemented in MEGA X (Kumar et al. 2018), with posterior manual edition. We performed BLAST searches in GenBank (http://blast.st-va.ncbi.nlm.nih.gov/Blast.cgi) in order to determine the approximate association of the obtained sequences with published records. Phylogenetic inference of Maximum Likelihood (ML), Maximum Parsimony (MP) and Neighbor Joining (NJ) was generated in MEGA X using Tamura and Nei’s (1993) model of nucleotide substitution and variable sites following a gamma distribution (TN93+G). Bayesian analysis was performed in Mr. Bayes v3.2.6 (Ronquist & Huelsenbeck 2003). Phylogenetic trees were later edited in FigTree v1.4.3 (Rambaut & Drummond 2012). We also conducted systematic searches in online academic databases (Google Scholar, Scielo, Scopus, Web of Science) for occurrence localities of *Cryptonanus agricolai*, in order to elaborate an updated distribution map of the species. We only considered as reliable those records that included a detailed description of how the specimen was identified.

## RESULTS

The individual UFES-MAM 3075 captured in a pitfall trap on 17th October 2015 was an adult male with evident scrotal testicles and dental age class 5 (all molars functional and permanent third premolar completely erupted; Tribe 1990). External, cranial and dental measurements are shown in Table 1. A combination of morphological aspects related to the upper premolars (P3 greater than P2) and fenestrae in the palate (long maxillopalatine fenestra, palatal fenestra, posterolateral foramen, incisive foramen and absent maxillary fenestra) led us to identify the specimen as belonging to the genus *Cryptonanus* (Fig. 1), according to didelphid identification key to genera (Rossi et al. 2012; Voss & Jansa 2009).

**Table 1.**
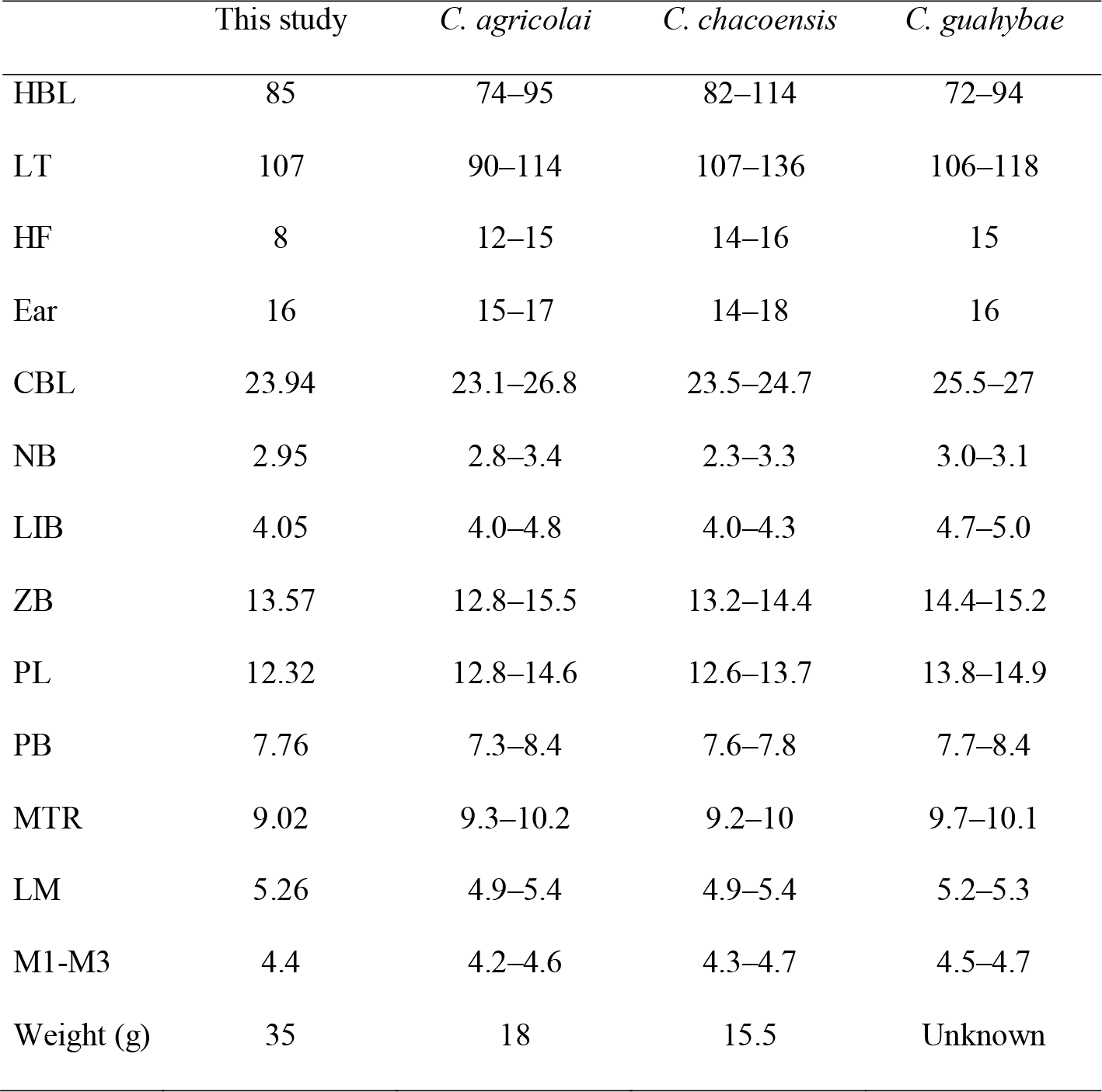
External, cranial and dental measurements of *Cryptonanus* species with occurrence in Brazil (Gardner 2008) and the study specimen (UFES-MAM 3075).

**Fig. 1.**
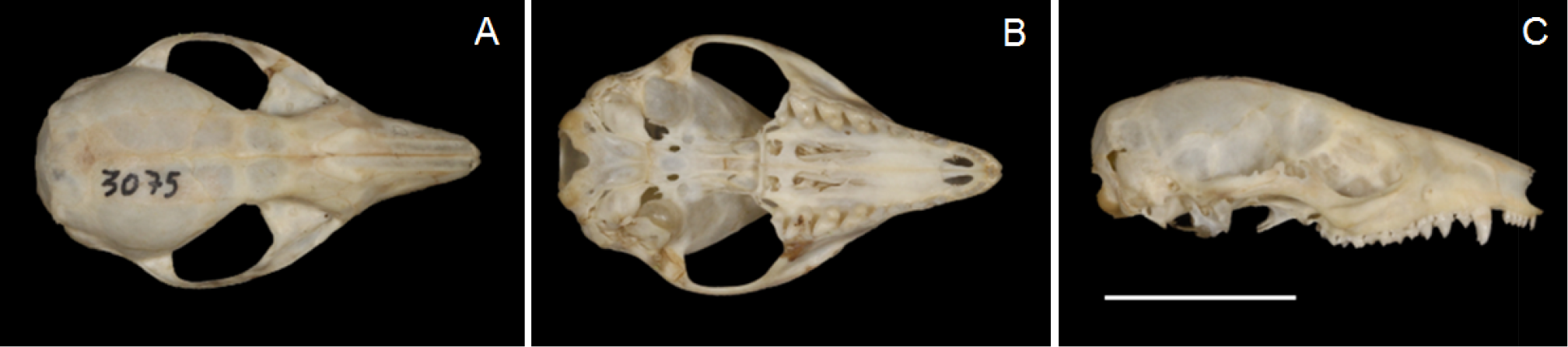
Dorsal, ventral, and lateral skull views of *Cryptonanus agricolai* from Pinheiros, state of Espírito Santo, Southeast Brazil (A–C; UFES-MAM 3075). Scale bar = 10 mm (Credit: Heitor Bissoli).

For accurate identification at the species level, we also used molecular approaches. In a BLAST search we found 98% identity with published sequences of *C. agricolai*. Through phylogenetic inference it was possible to identify the specimen as *C. agricolai*, since the sample of UFES-MAM 3075 was grouped with other previously identified specimens of the same species, in a highly supported clade (Bayesian posterior probability = 1). Maximum likelihood and Bayesian inference trees depicted the same topology - only the latter is here represented (Fig. 2). The sister status of *C. agricolai* and *C. chacoensis*, was also strongly supported, and these two taxa formed a marginally supported clade including *C. guahybae*, with *C. unduaviensis* as the outermost clade, then composing a monophyletic group for the genus *Cryptonanus*. Through systematic searches in online academic databases, it was possible to compile unambiguous records of *C. agricolai* in the literature (Table 2), and to estimate the current extent of occurrence (minimum convex polygon) for the species in 1 773 398 km^2^, (Fig. 3).

**Fig. 2.**
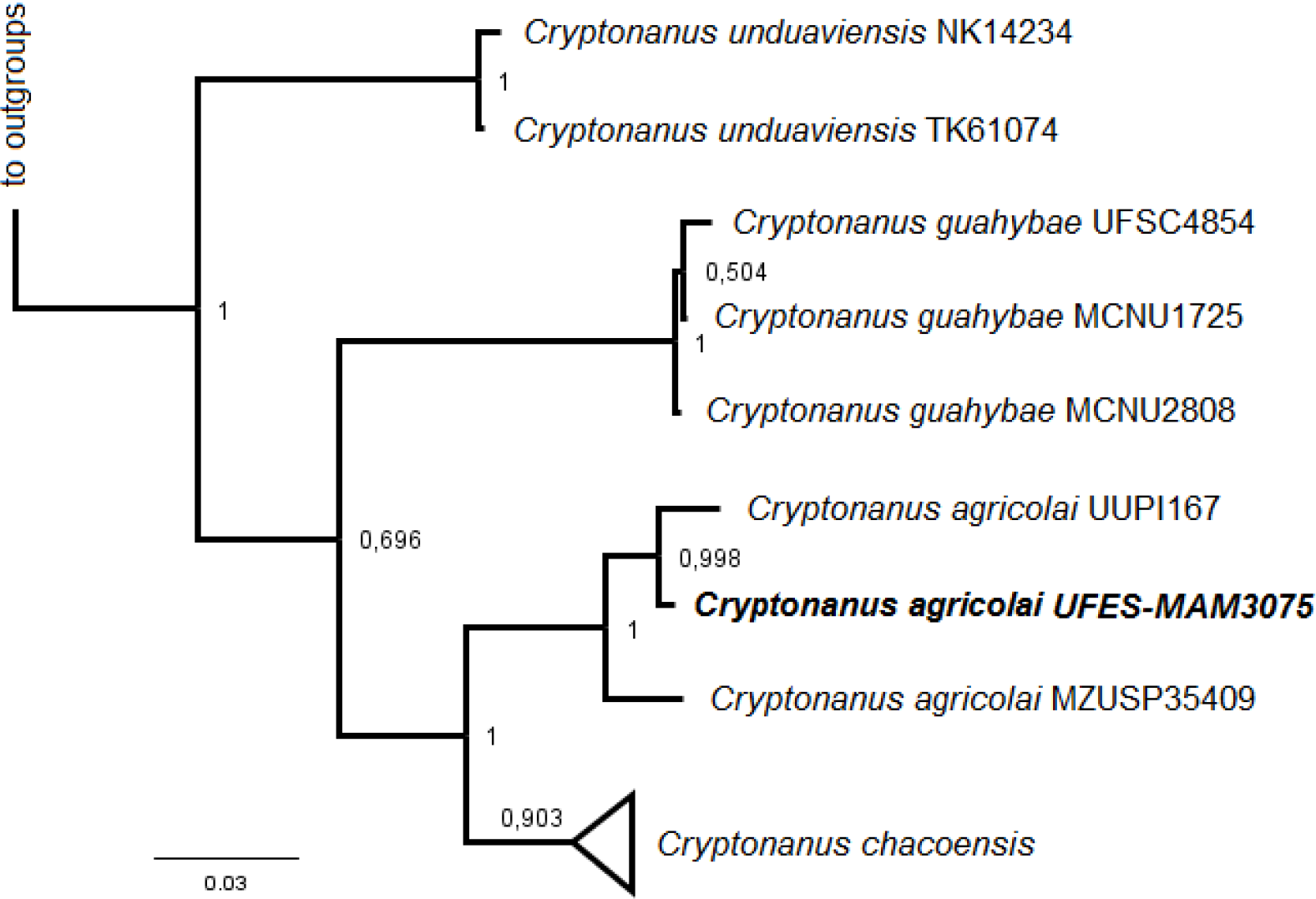
Phylogenetic inference of four *Cryptonanus* species based on Bayesian Analysis of 801 bp of mitochondrial Cytochrome B sequences, performed using the TN64 + G model. Values at the nodes refer to Bayesian posterior probabilities. In bold, the specimen reported in the present study (UFES-MAM 3075). *Gracilinanus peruanus* and *Thylamys citellus* were used as outgroups.

**Table 2.**
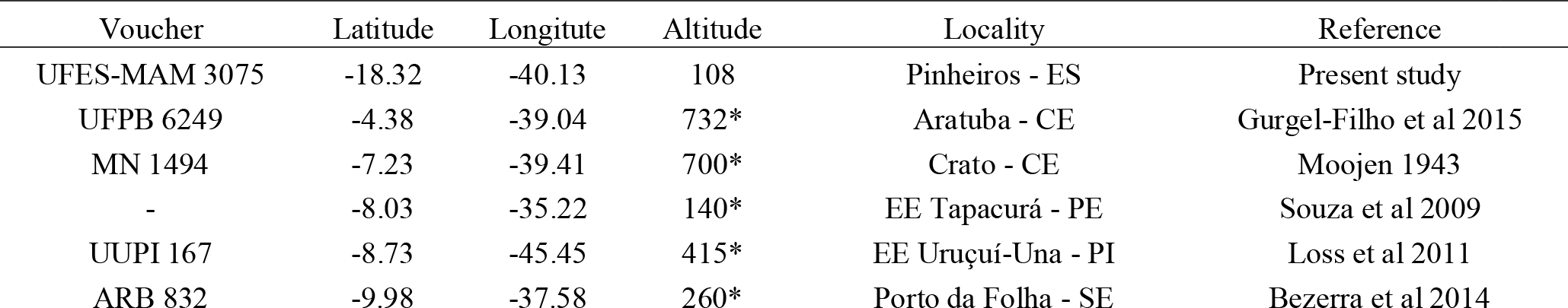
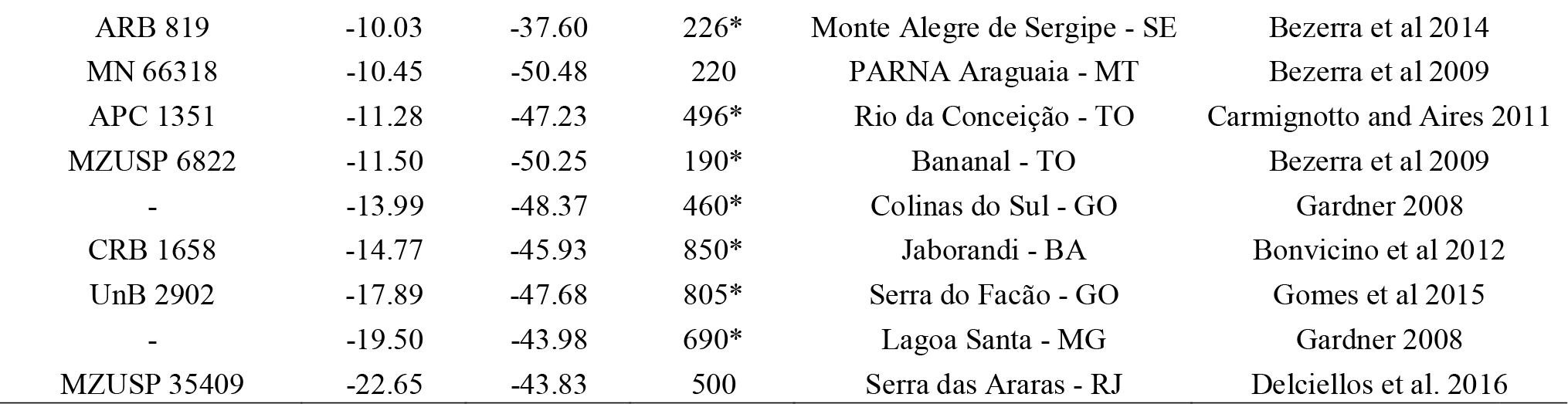
Localities of *Cryptonanus agricolai* obtained from the literature and used as points to elaborate the map of extension of occurrence for the species. Acronyms after hyphens indicate Brazilian states; BA = Bahia, CE = Ceará, ES = Espírito Santo, GO = Goiás, MG = Minas Gerais, MT = Mato Grosso, PE = Pernambuco, PI = Piauí, RJ = Rio de Janeiro, SE = Sergipe, TO = Tocantins. Asterisks (*) indicate values inferred from the geographic coordinates provided by authors.

| Voucher | Latitude | Longitude | Altitude | Locality | Reference |
| --- | --- | --- | --- | --- | --- |
| UFES-MAM 3075 | -18.32 | -40.13 | 108 | Pinheiros - ES | Present study |
| UFPB 6249 | -4.38 | -39.04 | 732* | Aratuba - CE | Gurgel-Filho et al 2015 |
| MN 1494 | -7.23 | -39.41 | 700* | Crato - CE | Moojen 1943 |
| - | -8.03 | -35.22 | 140* | EE Tapacurá - PE | Souza et al 2009 |
| UUPI 167 | -8.73 | -45.45 | 415* | EE Uruçuí-Una - PI | Loss et al 2011 |
| ARB 832 | -9.98 | -37.58 | 260* | Porto da Folha - SE | Bezerra et al 2014 |
| ARB 819 | -10.03 | -37.60 | 226* | Monte Alegre de Sergipe - SE | Bezerra et al 2014 |
| MN 66318 | -10.45 | -50.48 | 220 | PARNA Araguaia - MT | Bezerra et al 2009 |
| APC 1351 | -11.28 | -47.23 | 496* | Rio da Conceição - TO | Carmignotto and Aires 2011 |
| MZUSP 6822 | -11.50 | -50.25 | 190* | Bananal - TO | Bezerra et al 2009 |
| - | -13.99 | -48.37 | 460* | Colinas do Sul - GO | Gardner 2008 |
| CRB 1658 | -14.77 | -45.93 | 850* | Jaborandi - BA | Bonvicino et al 2012 |
| UnB 2902 | -17.89 | -47.68 | 805* | Serra do Facão - GO | Gomes et al 2015 |
| - | -19.50 | -43.98 | 690* | Lagoa Santa - MG | Gardner 2008 |
| MZUSP 35409 | -22.65 | -43.83 | 500 | Serra das Araras - RJ | Delciellos et al. 2016 |

**Fig. 3.**
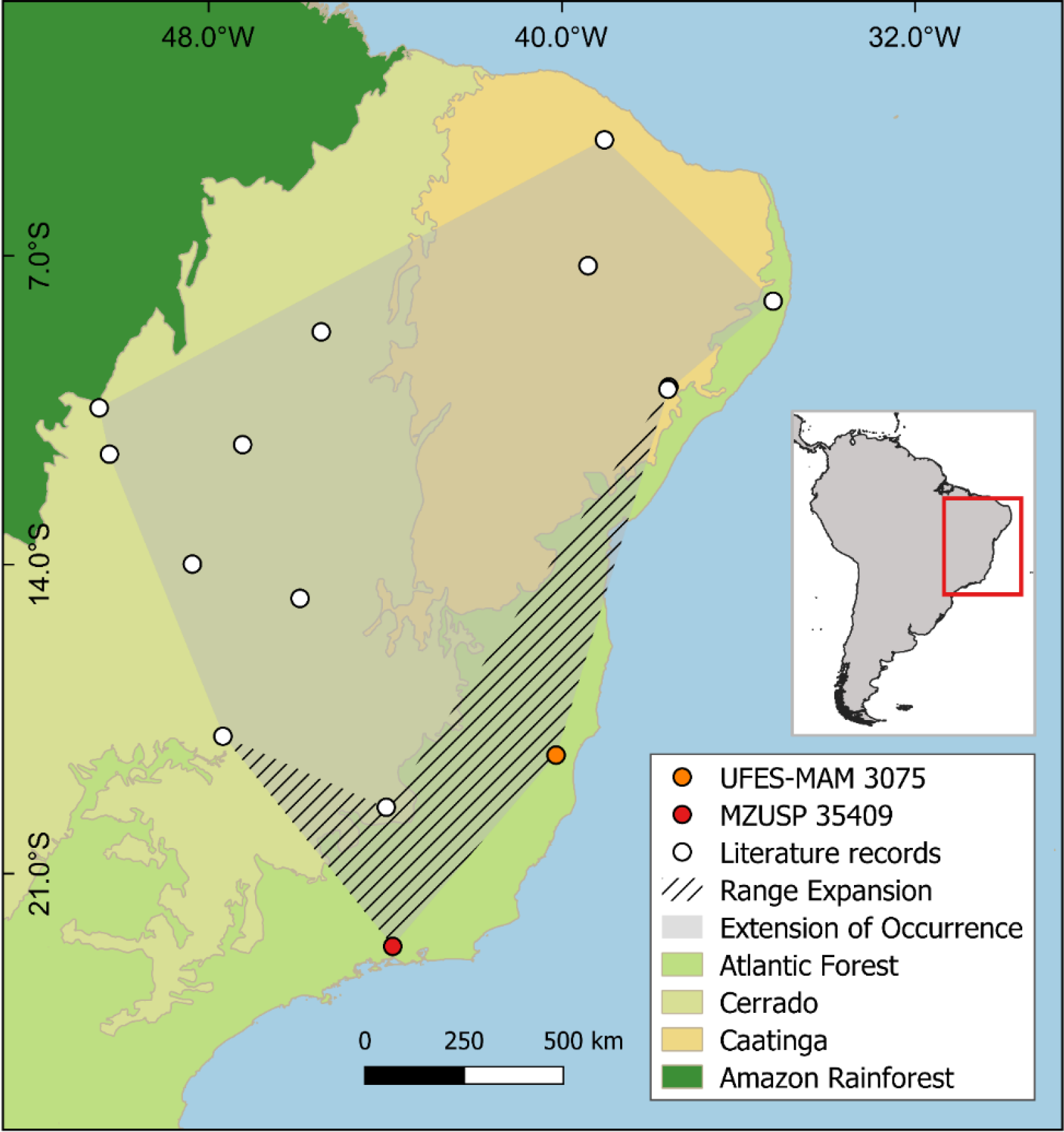
Updated extent of occurrence and range expansion of *Cryptonanus agricolai* to the Atlantic Forest, according to the new records.

## DISCUSSION

The specimen here identified as *Cryptonanus agricolai* represent the first record of the species in the Central Atlantic Forest Corridor (Brazil 2006; Camara & Galindo-Leal 2009), a region formerly considered outside its distributional range (Carmignotto et al. 2016). Previously, two records of *Cryptonanus* had already been registered in the Atlantic Forest; however, in both cases the authors did not identify the specimens at species level (see Souza et al. 2010; Delciellos et al. 2016). Therefore, the present study expands the extent of occurrence of the species in 324 452 km^2^ (Fig. 3; hatched area), updating to 1 773 398 km^2^ the total area of occurrence. Additionally, our record from Pinheiros, state of Espírito Santo, set the lower altitudinal limit for *C. agricolai* at 108 m above sea level (Table 2). Moreover, according to the geographical coordinates of a trap site mentioned in Bonvicino et al. (2012) we inferred what can be recognized as the highest altitudinal limit of the species: 850 m; an increase of almost a 100 m of that previously suggested (400 to 760 m; Gardner 2008). Hence, we can now establish the altitudinal range of *C. agricolai* from 108 to 850m.

Besides changes in distributional and altitudinal ranges of *C. agricolai*, our work confirms the occurrence of the species, often associated with xeric habitats and open vegetation, in a central region of the Atlantic Forest domain, with points of occurrence far from the contact zones of this biome with either the Cerrado or Caatinga domains. Although the individual was captured in an ecoregion distinct from that usually related to *C. agricolai*, the species can still continue to be associated with xeric-open habitats, since the sampling site in Reserva Biológica Córrego do Veado is dominated by patches of vegetation called Mussununga Such formations are characterized by phytophysiognomies ranging from grasslands to woodlands over sandy soils that have high water retention, with a consistently hard and impermeable cementation layer, which causes flooding stress in the rainy season and drought stress in the dry season (Mecke et al. 2002; Horbe et al. 2004; Saporetti-Junior et al. 2012). Despite the fact that Mussununga formations are threatened by many factors (e.g. fire, logging, road construction, and biological invasion), such kind of vegetation is still underrepresented in the context of studies conducted in the Brazilian Atlantic Forest (Eisenlohr et al. 2015; Heringer et al. 2019).

In addition to *C. agricolai*, another species of the genus occurs in the South America diagonal of open formations (e.g. Chaco, Cerrado and Caatinga domains; Prado & Gibbs 1993): *C. chacoensis*, which is widely distributed in Gran Chaco, but there is also records for this species in southwest areas of the Cerrado Domain (Teta & Días-Nieto 2019). Since there are strong evidence that each species may constitute a species complex, pending a formal revision (Astúa 2015), and considering the possibility of sympatry between *C. agricolai* and *C. chacoensis* in the southwest portion of Cerrado, as well as the morphological similarities shared by them, we did not include in the updated occurrence map (Fig. 3) records of *C. agricolai* provided by articles that report its occurrence in this area, but without detailed description of identification diagnosis (Caceres et al. 2008; Martin et al. 2012; Hannibal & Neves-Godoi 2015; Gonçalves et al. 2018). Although delimiting the western limits of *C. agricolai* is somewhat outside the scope of our work, we emphasize the importance of meticulous and comprehensive identifications, as well as descriptions of how species were diagnosed when distribution records are given, especially in areas with possible sympatry of cryptic genus and species.

When the genus was described, authors mentioned that significant range extensions of *Cryptonanus* could be expected by surveys in extralimital savanna landscapes, specially by pitfall trapping (Voss et al. 2005; Dias et al. 2015). In the Central Atlantic Forest Corridor (Brazil 2006; Camara & Galindo-Leal 2009), small mammal surveys using pitfall traps are still scarce (Bovendorp et al. 2017), despite their obvious and demonstrated efficiency to, trap elusive species (Rocha et al. 2015). Protocols that include this and other alternative kinds of trap methods could contribute largely to fill in many gaps on species distribution, as showed here. The records presented herein as *C. agricolai* expanded the geographic distribution of the species approximately 430 km eastward (UFES-MAM 3075; Fig. 3) and 360 km southward (the recognition of MZUSP 35409 as *C. agricolai*), throughout an area of 324 452 km^2^ (hatched area; Fig. 3). This augment corresponds to a 22% of increase in the extent of occurrence previously known for the species and indicates that *C. agricolai* distribution is significantly larger, extending over a substantial portion of the Atlantic Forest ecoregion.

## ACKNOWLEDGMENTS

We would like to thank Elisandra Chiquito and Roger Guimarães for reading and commenting on earlier versions of this manuscript. Luciana Conde and Roberta Paresque for fieldwork coordination and support; João Luiz Guedes and Yuri Leite for assisting in laboratory techniques and molecular analysis. EBG had no funding while working on this manuscript; LPC has received continuous support from Fundação de Amparo à Pesquisa e Inovação do Espírito Santo (FAPES), Coordenação de Aperfeiçoamento de Pessoal de Nível Superior (CAPES) and Conselho Nacional de Desenvolvimento Cientifico e Tecnológico (CNPq).

